# Evolutionary responses of energy metabolism, development, and reproduction to artificial selection for increasing heat tolerance in *Drosophila subobscura*

**DOI:** 10.1101/2022.02.03.479001

**Authors:** Andrés Mesas, Luis E. Castañeda

## Abstract

Adaptations to warming conditions exhibited by ectotherms include increasing heat tolerance but also metabolic changes to reduce maintenance costs (metabolic depression), which can allow them to redistribute the energy surplus to biological functions close to fitness. Currently, there is evidence that energy metabolism evolves in response to warming conditions but we know little about how the rate of temperature change during heat stress determines the evolutionary response of metabolism and the consequences on life-history traits. Here, we evaluated the evolutionary response of energy metabolism (metabolic rate and activity of enzymes of the glucose-6-phosphate branchpoint) and life-history traits to artificial selection for increasing heat tolerance in *Drosophila subobscura*, using two different thermal selection protocols for heat tolerance: slow and fast ramping protocols. We found that the increase in heat thermal tolerance was associated with a reduction of the hexokinase activity in the slow-ramping selected lines, and a slight reduction of the glucose-6-phosphate dehydrogenase activity in the fast-ramping selected lines. We also found that the evolution of increased heat tolerance increased the early fecundity in selected lines and increased the egg-to-adult viability only in the slow-ramping selected lines. However, heat tolerance evolution was not associated with changes in the metabolic rate in selected populations. This work shows heat tolerance can evolve under different thermal scenarios but with different evolutionary outcomes on associated traits depending on the intensity of thermal stress. Therefore, spatial and temporal variability of thermal stress intensity should be taken into account to understand and predict the adaptive response to ongoing and future climatic conditions.

## Introduction

Environmental thermal stress involves strong selective pressures that can lead to important evolutionary changes depending on genetic variation exhibited by natural populations (Umina et al., 2005; Crozier & Hutchings, 2014; Chick et al., 2021). The increasing environmental temperatures cause fitness declines and consequently changes in abundance and distribution of ectotherm populations (Deutsch et al., 2008; Angilletta, 2009; Chen et al., 2011). Therefore, knowing and exploring how ectotherms will withstand global warming is crucial.

The ability to tolerate high temperatures implies that organisms must exhibit an array of responses at different levels of organization, allowing them to maintain their biological performance under these conditions (Huey & Kingsolver, 1993; Pörtner et al. 2006; Mesas et al. 2021; McGaughran et al. 2021). Among these responses, metabolic depression, defined as a reduction of the energy expenditure associated with the maintenance of organisms, is considered an important physiological mechanism related to resistance to environmental stress (Guppy & Withers, 1999; Storey & Storey, 2004). Plastic and evolutionary changes in energy metabolism should have important ecological and evolutionary consequences when organisms are exposed to stressful conditions because it should allow them to reduce their minimum energetic requirements and redistribute the energy surplus to biological functions close to fitness (Atkinson, 1977; Storey & Storey, 2004). For example, individuals with a low standard metabolic rate (SMR) have higher survival compared to those individuals with high SMR (Artacho & Nespolo, 2009). Additionally, the evolution of increased resistance to desiccation, starvation, hypoxia, and high temperatures has been also accompanied by a reduction in metabolic rate (Hoffmann & Parson, 1989; Padfield et al. 2016; Regan et al. 2017; Brown et al. 2019; but see Djawdan et al. 1997), suggesting that energy-saving phenotypes are favored under stressful conditions.

On a broader scale, metabolic depression as a physiological response to high temperature could explain the countergradient variation in metabolic rate proposed in the Metabolic Cold Adaptation hypothesis (Addo-Bediako, 2002; Gaston et al. 2009), which establishes that ectothermic species from temperate habitats have lower metabolic rates than those living in colder environments (Addo-Bediako et al. 2002; Sylvestre et al. 2007; Schaefer & Walters 2010; Sinnatamby et al. 2015; Pilakouta et al. 2020). Also, metabolic depression has represented an adaptive response in phytoplankton populations, which evolved in warm environmental conditions and showed an increase in their thermal tolerance accompanied by a reduced metabolic rate (Padfield et al. 2016). However, there is also evidence that *Drosophila simulans* showed an increase in metabolic rate after 60 generations of evolution in a hot environment (Mallard et al. 2018). Regardless of the direction of the metabolic response to warm conditions, these results indicate that metabolism can evolve when populations are exposed to stable warm conditions. However, under natural conditions, organisms are exposed to fluctuating environmental conditions, which can result in different evolutionary outcomes (Colinet et al 2015). For instance, Mesas et al (2021) found an evolutionary increase in heat tolerance in *D. subobscura* regardless of the ramping rate used during thermal assays, but changes in the thermal performance curve of the evolved populations did depend on the ramping rates used during the selection. This evidence suggests that heat tolerance may evolve in response to global warming, but responses of associated traits, including metabolic rate, would depend on the rate of temperature change during heat stress (Santos et al 2012). Indeed, individuals with slow metabolism are expected to have a higher heat tolerance than individuals with fast metabolism because the latter depletes energy resources faster than the former during heat stress (Rezende et al. 2011; Santos et al. 2012).

Metabolism is an emergent property of interactions between different cellular processes and metabolic pathways dependent on temperature (Berrigan & Hoang, 1999; Montooth et al. 2003; Brown et al. 2004; Matoo et al. 2019). Then, selection under heat stress should involve evolutionary changes not only in metabolic rate but also in the underlying metabolic pathways. For example, several genes associated with the glucose-6-phosphate (G6P) branchpoint, an important pathway in the storage and use of molecules such as glucose, glycogen, triglycerides, and ATP, have shown clinal variation in their allelic frequencies along latitude (Oakeshott et al 1983; Duvernell & Eanes, 2000; Verrelli & Eanes, 2001; Rank & Dahlhoff, 2002, Sezgin et al. 2004). These findings suggest that the functional and structural properties of the enzymes associated with energy metabolism are under selection, likely driven by environmental temperature. Additionally, genetic variation in the G6P branchpoint results in variability in the metabolic flux in ectotherms (Verrelli & Eanes, 2001), facilitating evolutionary responses among fitness-related traits under stressful conditions (Zera & Zhao, 2003). Evidence from a laboratory selection experiment suggests that the evolution under warm conditions results in a down-regulation of metabolic pathways and an increase in fecundity in *D. simulans* (Mallard et al. 2018). Thus, adaptation to warming environments should involve metabolic readjustments to reduce negative effects on fitness-related traits such as reproduction and development traits (Sales et al., 2018; Parratt et al., 2021; García-Robledo & Baer, 2021).

In this work, we evaluated the evolutionary response of energy metabolism and life-history traits to artificial selection for increasing heat tolerance in *D. subobscura*, using two different thermal selection protocols for heat tolerance: slow and fast ramping protocols. According to the assumption that individuals with low metabolic rates can withstand longer heat stress, we hypothesized that increased heat tolerance evolves with a reduction of energy metabolism, especially if the thermal selection is performed using a long (slow-ramping rate) versus short (fast-ramping rate) protocols to estimate heat tolerance. Additionally, we proposed that individuals with low metabolic rates will have an energy surplus, which should have consequences on fitness-related traits (e.g., high developmental and reproductive success). To evaluate these hypotheses, we measured the routine metabolic rate (RMR) and the activity of enzymes associated with the G6P branchpoint as proxies of the energy metabolism in evolved populations of *D. subobscura* previously selected for increasing heat tolerance (Mesas et al. 2021). We measured the activity of these enzymes in flies exposed to two different thermal conditions (non-stress: 21 °C, and stress: 32 °C) to evaluate the thermal plasticity of G6P-related enzymes. Additionally, we evaluated the early fecundity and egg-adult viability in the control and selected populations to determine the effects of heat tolerance evolution on the life-history traits of *D. subobscura*.

## Methods

### Heat knockdown temperature selection

A mass-bred population of *D. subobscura* was established from the offspring of 100 isofemale lines derived from females collected in Valdivia, Chile (39.8 °S 73.2 °W). Specifically, we collected inseminated females using banana-yeast traps and these females were individually placed into vials with David’s killed-yeast *Drosophila* medium (hereafter *Drosophila* medium) to establish isofemale lines. At the next generation, 10 females and 10 males from each one of the 100 isofemale lines were transferred to an acrylic cage (27 × 21 × 16 cm3) to set up one large outbred population (>1500 breeding adults). In the next generation, eggs were collected and transferred to 150 mL bottles with fly food at a density of 150 eggs/bottle. The bottles were divided into 3 groups (15 bottles/group) and the flies that emerged from each group were transferred to one acrylic cage, resulting in a total of three population cages: R1, R2, and R3. After three generations, when the population cages reached a large population size and environmental effects were removed, each replicated cage (R1, R2, and R3) was divided into four population cages. Thus, our experimental design had 12 population cages with more than 1000 individuals each, which were assigned to four different artificial selection protocols in triplicate: fast-ramping selection, fast-ramping control, slow-ramping selection, and slow-ramping control lines (Fig. S1). During all generations, the population cages were maintained at 21 °C (12:12 light:dark cycle) and fed with *Drosophila* medium in Petri dishes before the artificial selection experiment.

For each replicate line, we randomly took 160 four-day-old virgin females, which were individually placed in a vial with *Drosophila* medium together with two males from the same replicate line. After two days of mating, males were discarded from each vial and the presence of eggs or first-instar larvae was checked. Of these 160 mated females, 120 females were randomly chosen to measure their knockdown temperature and the remaining females were discarded. These 120 females were individually placed in a capped 5-mL glass vial, which was attached to two racks, each one with a capacity to attach 60-capped vials. Racks were immersed in a water tank with an initial temperature of 28 °C, which was controlled by a heating unit (Model ED, Julabo Labortechnik, Seelbach, Germany). After an equilibration time of 10 min, the temperature was increased at a rate of 0.08 °C min-1 for the slow-ramping selection protocol, or 0.4 °C min-1 for the fast-ramping selection protocol. Thermal assays ended when the temperature reached 45 °C, the temperature at which all females collapsed. Each assay was video recorded with a high-resolution camera (D5100, Nikon, Tokyo, Japan). Videos were visualized to score the knockdown temperature for each female fly, which was defined as the temperature at which each fly ceased to move. For the selection, we selected vials containing the offspring of the 40 most tolerant flies (upper 30% of each assay) to found the next generation (offspring was never exposed to heat stress). Specifically, from the offspring of the selected females (40) we collected four virgin females to re-established the original number of 160 females. For the fast-control and slow-control lines, we measured the heat tolerance of 40 females as mentioned above and randomly selected the offspring of 10 of them to found the next generation.

Artificial selection for heat tolerance was performed for 16 generations, after which flies were dumped in acrylic cages and maintained without selection for heat tolerance at 21°C (12:12 light-dark cycle) until the measurement of metabolic and life-history traits. For logistical reasons, we used the fast-ramping control experimental line as the control line for statistical comparisons.

### Metabolic rate

The routine metabolic rate (RMR) was measured as the amount of CO_2_ produced by individual four-day-old virgin females from generation 26. RMR was measured in 15 females from each replicate line, reaching a total of 135 flies (3 thermal selection protocols x 3 replicate lines x 15 flies). Before metabolic measurements, flies were ice-cold anesthetized and weighted using a microbalance (MXA5, Radwag, Czech Republic). Then, flies were individually placed in a metabolic chamber, which was made of 2 cm of a Bev-A-Line tube. The metabolic chambers were connected to an eight-channel multiplexer system (RM8, Sable Systems International, NV, USA) and a 5 mm^2^ fabric mesh was placed between the connectors to avoid flies passing from the metabolic chambers to the respirometry system. This system included seven chambers containing individual flies and one empty chamber used for baseline. Metabolic chambers were placed in dark conditions within a thermal cabinet (PC-1, Sable Systems International, NV, USA) at a temperature of 21 °C, which was controlled by a temperature controller (PELT-5, Sable Systems International, NV, USA). CO_2_-free air was pumped into chambers at a flow of 15 mL/min (Sierra MFC-2, Sable Systems International, NV, USA) and after 30 min of equilibration, CO_2_ production was sequentially measured in each chamber for 15 min using an infrared CO_2_ analyzer with a resolution of 1 ppm (Li-6251, LI-COR Bioscience, NE, USA). Measurements of CO_2_ were recorded, transformed to metabolic rate (μl CO_2_/h), and analyzed using the software Expedata (Sable Systems International, NV, USA).

### Enzyme activity

The catabolic activities of enzymes from the G6P branchpoint were measured in four-day-old females from generation 27 through the appearance of NADPH in kinetic assays (Montooth et al., 2003). We evaluated the activity of hexokinase (HEX: E.C. 2.7.1.1) involved in glucose metabolism, phosphoglucomutase (PGM; E.C. 5.4.2.2) involved in glycogen metabolism, phosphoglucose isomerase (PGI; E.C. 5.3.1.9) involved in ATP metabolism; and glucose-6-phosphate dehydrogenase (G6PD; E.C. 1.1.1.49) involved in lipids metabolism.

For each replicate line, 100 females were collected from each population cage and pooled in groups of 10 flies each. We measured the enzyme activity of the G6P branchpoint in flies exposed to two thermal conditions: thermal-stressed flies (five replicates were exposed to 32 °C per 1 h and then, flies were allowed to recover at 21 °C for 2 h); and non-stressed flies (five replicates were maintained at 21 °C for 3 h). Then, flies were quickly frozen in liquid nitrogen, homogenized in 1 mL of homogenization buffer (0.01 M KH2PO4, 179 1.0 mM EDTA, pH 7.4), and centrifuged at 2,000 rpm for 2 min at 4 °C. The supernatant was aliquoted and maintained at −80 °C until enzymatic assays. The buffers for each assayed enzyme were as follow. **HEX:** 20 mM TrisHCl, 0.5 mM NADP, 0.2 mM MgCl_2_, 0.36 mM ATP, 5 mM D-glucose, 0.23 units/mL G6PD, pH 8.0. **PGI:** 20 mM TrisHCl, 0.28 mM NADP, 3 mM fructose-6-phosphate, 1.37 units/mL G6PD, pH 8.0. **PGM:** 20 mM TrisHCl, 0.5 mM NADP, 1 mM MgCl_2_, 0.83 mM glucose-1-phosphate, 3.1 units/mL G6PD, pH 8.0. **G6PD:** 20 mM TrisHCl, 0.2 mM NADP, 18.8 mM MgCl_2_, 3.5 mM glucose-6-phosphate, pH 8.0. Enzymatic assays were performed by adding 25 μL extracts of flies with a protein concentration of 3500 μg/mL and 200 μL of respectively assay buffer in ultraviolet plates with flat bottom wells. Immediately, changes in optical density (OD) at a wavelength of 340 nm was measured with multiple reads per well in a microplate reader (Infinite 200 Pro, Tecan) previously heated to 29 °C. The OD was measured each 25 s during 10 min to HEX and PGI, every 10 s during 10 minutes to PGM, and every 45 s during 15 min to G6PD. Enzymatic activities were estimated in triplicate and mean OD was analyzed. Blank wells containing buffer assays and double-distilled water (instead of fly extracts) were measured to establish the basal optical density of each reaction. Finally, the enzymatic activity was calculated as the change in optical density over time.

### Early fecundity and egg-to-adult viability

Early fecundity was measured in females from generation 26. For each replicate line, we collected 10 virgin females, which were individually placed in vials with *Drosophila* medium together with two unrelated males from the same replicate line. After 24 h of mating, females were transferred to new vials and the number of oviposited eggs in each vial was counted daily using a stereomicroscope. This procedure was repeated every 24 h for eight days to obtain the accumulated fecundity for each female. Additionally, we measured the egg-adult viability as follows. Vials containing eggs oviposited by 5-day-old females were retained because *D. subobscura* reaches its peak fecundity between the age of 3 and 7 days (Foucaud et al., 2016). For each vial, 20 eggs were randomly transferred to a new vial and the number of emerging adults was counted.

### Statistical analysis

Normality and homoscedasticity were tested for all variables, and only the enzyme activity of HEX was squared-root transformed to meet the parametric assumptions. Then, a mixed linear model (hereafter ‘full model’) was used to evaluate the effects of thermal selection (fixed effect) and replicate lines nested within thermal selection (random effect) on heat knockdown temperature (eq. 1), body mass (eq. 2) and RMR (including body mass as covariate) (eq. 3):

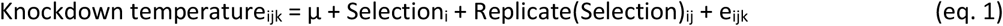

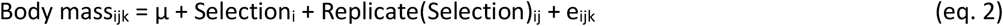

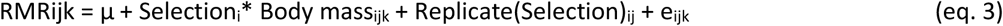

For the enzyme activities, the full model included the interaction between the thermal exposure and thermal selection as fixed effects (eq. 4):

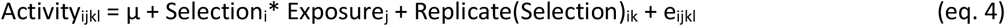

Early fecundity was analyzed using a generalized linear model with a Poisson (link = log) distribution with thermal selection, oviposition days and their interaction were considered as fixed effects, while replicate lines nested within thermal selection were considered as a random effect (eq. 5). As fecundity was measured for each female along 8 days, females’ identity (ID) was included in the model to accomplish for the repeated-measures design:

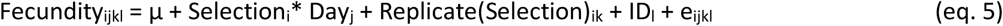

Finally, egg-to-adult viability was estimated as the proportion of emerging flies per vial, considering that 20 eggs were placed in a vial. This proportion was analyzed using a generalized linear model with a binomial distribution with thermal selection as fixed effect, and replicate lines nested within thermal selection were considered as a random effect (eq. 6):

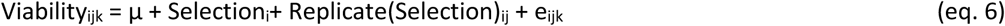

All statistical analysis was performed in R (R Core Team, 2021) using the function lmer and glmer of the lme4 package for the generalized mixed model (Bates et al. 2014). Post-hoc analyses were performed using a Tukey-adjusted method using the emmeans package (Lenth, 2021). All data and scripts are available online (Mesas and Castañeda, 2022; https://doi.org/10.6084/m9.figshare.20373180.v1).

## Results

### Knockdown temperature evolution

To test the evolutionary response of knockdown temperature to selection for higher thermal tolerance, we compared the knockdown temperature for control and selected lines in the generation 25. This comparison was independently performed for each thermal selection protocol because thermal tolerance is always higher in fast ramping assays estimate than in slow ramping assays (Rezende et al. 2011; Castañeda et al. 2015). We found that fast-ramping selected lines evolved for significantly higher knockdown temperatures than fast-ramping control lines (mean_fast-ramping_ ± SD = 38.59 ± 0.98 °C and mean_fast-control_ ± SD = 37.61 ± 1.45 °C; F_1,4_ = 56.4, P = 0.002). Similarly, slow-ramping selected lines evolved significantly higher knockdown temperatures than slow-ramping control lines (mean_slow-ramping_ ± SD = 35.92 ± 0.67 °C and mean_slow-control_ ± SD = 35.14 ± 0.81 °C; F_1,4_ = 83.5, P = 0.001). These results show that differences in heat tolerance between selected and control lines were maintained after nine generations without selection for heat tolerance (Mesas et al. 2021).

### RMR and body mass

Body mass was not significantly different between the control and selected lines (*X*^2^_2_ = 0.46, P = 0.794, Fig. 1A) and it did not differ between replicate lines (*X*^2^_1_ = 0.03, P = 0.862). On the other hand, as expected, RMR and body mass were significantly correlated (r = 0.29, t_132_ = 3.48, P = 0.0007, Fig. 1B). Then, body mass was included as covariate in the mixed linear model to test the effects on RMR, which showed that RMR did not differ between thermal selection regimens (*X*^2^_2_ = 0.28, P = 0.870; Fig. 1C) and neither between the replicate lines (*X*^2^_1_ = 0, P = 1).

**Figure 1.**
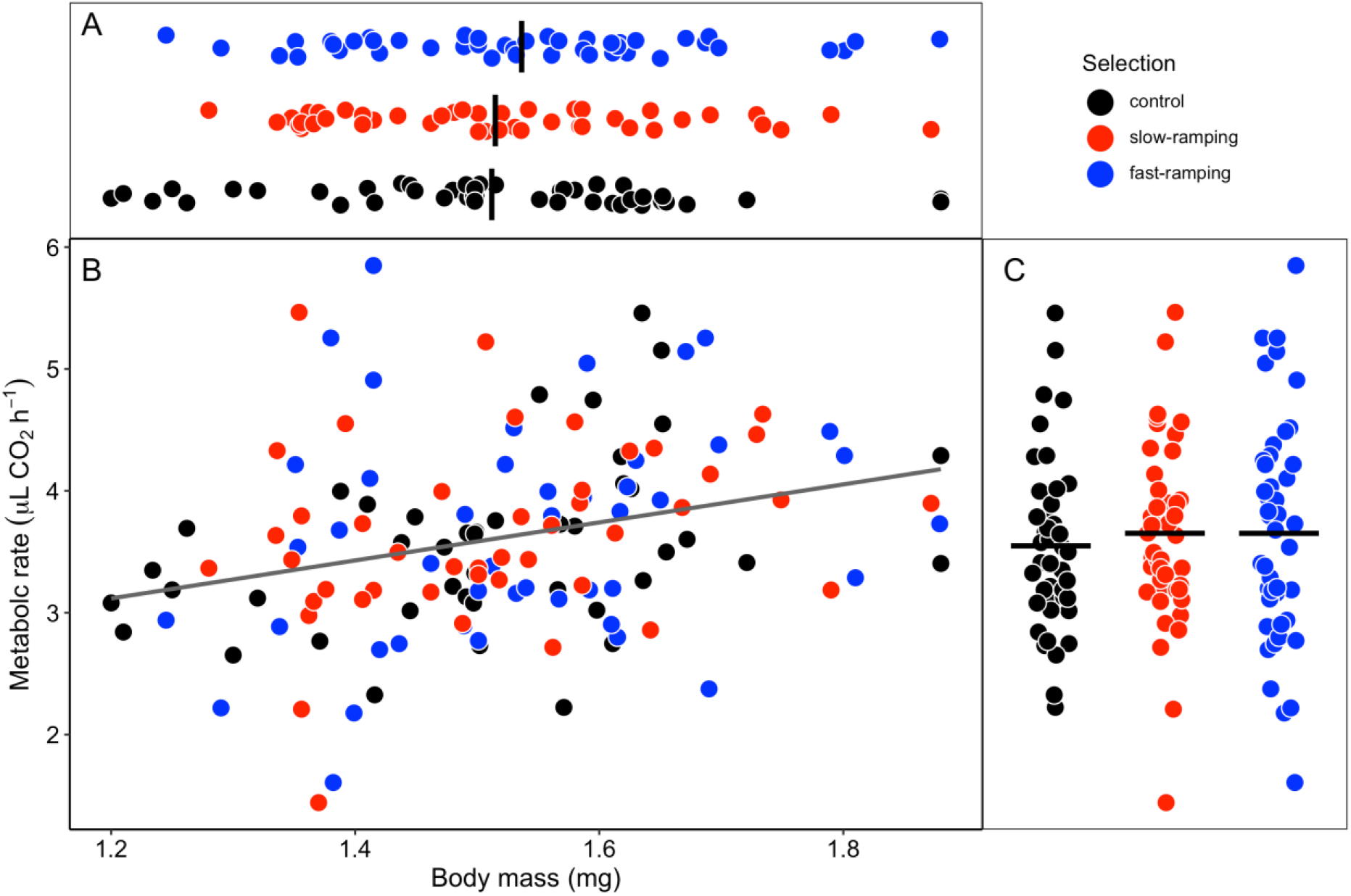
Body mass (A), the relationship between body mass and metabolic rate (B), and resting metabolic rate (C) for flies of control lines (black circles), slow-ramping selected lines (red circles), and fast-ramping selected lines (blue circles) for increasing heat tolerance in *Drosophila subobscura*. In panel B, the grey line represents the linear regression between both traits. Non-significant differences between thermal selection regimens were found for body mass and resting metabolic rate (see text).

### Enzyme activity

For HEX activity (Fig. 2A), we found a significant effect of the thermal selection regimens (*X*^2^_2_ = 9.32, P = 0.009), a significant decrease of HEX activity after heat stress (*X*^2^_1_ = 4.46, P = 0.035), and a non-significant interaction between thermal selection and thermal exposure (*X*^2^_2_ = 2.28, P = 0.320). Looking for selection effects on each exposure treatment (Fig. 2A), we found that HEX basal activity evolved to lower values in the slow-ramping selected lines (Tukey P-adjusted value = 0.02), whereas fast-ramping selected lines showed similar HEX basal activity as control lines (Tukey P-adjusted value = 0.35). On the other hand, the induced activity of HEX was not different between thermal selection lines (Tukey P-adjusted value > 0.5). For PGI activity (Fig. 2B), we did not find significant effects of thermal selection (*X*^2^_2_ = 3.43, P = 0.180), thermal exposure (*X*^2^_1_ = 0.42, P = 0.517), neither significant interaction between these factors (*X*^2^_2_ = 2.10, P = 0.349). For PGM activity (Fig. 2C), we did not find significant effect of the thermal selection regimens (*X*^2^_2_ = 0.70, P = 0.706), thermal exposure (*X*^2^_1_ = 1.42, P = 0.233), neither significant interaction between these factors (*X*^2^_2_ = 1.16, P = 0.560). Finally, G6PD activity was significantly different between the thermal selection regimens (*X*^2^_2_ = 6.7, P = 0.035; Fig. 2D). On the other hand, G6PD activity did not show significant differences between thermal exposure (*X*^2^_1_ = 1.35, P = 0.245) neither a significant interaction between thermal selection and thermal exposure (*X*^2^_2_ = 1.78, P = 0.411). Looking for selection effects on each exposure treatment, we did not find a posteriori significant differences between thermal selection regimens within each thermal stress treatment (Tukey P-adjusted value > 0.05), probably because differences between thermal selection regimens were close to the significance threshold (α = 0.05).

**Figure 2.**
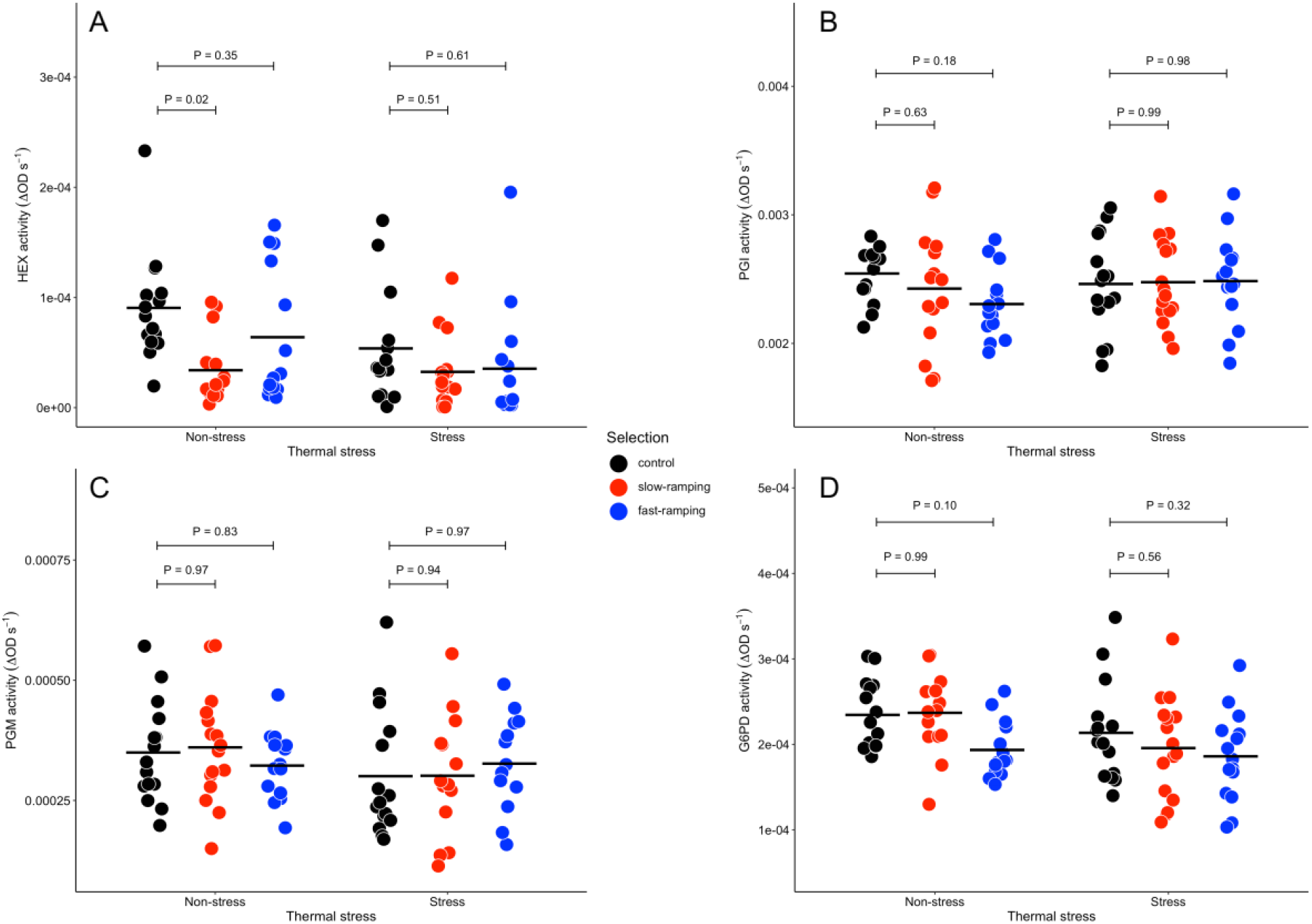
The activity of enzymes related to G6P branchpoint of flies of control lines (black circles), slow-ramping selected lines (red circles), and fast-ramping selected lines (blue circles) for increasing heat tolerance in *Drosophila subobscura* exposed to non-stressful (21 ºC) and stressful conditions (32 ºC): (A) hexokinase (HEX); (B) phosphoglucoisomerase (PGI); (C) phophoglucomutase (PGM); and (D) glucose-6-phosphate dehydrogenase (G6PD). Horizontal lines show the mean for each group. P-values above whiskers show the results of a posteriori comparisons between control and selected lines using Tukey tests.

Regarding the variability present among replicate lines, we found that the replicate lines showed a non-significant effect on the activity of all studied enzymes (*X*^2^_1_ = 0, P = 1).

### Early fecundity and egg-to-adult viability

Cumulative fecundity (Fig. 3) showed a significant effect of thermal selection (*X*^2^_2_ = 17.83, P = 0.0001), oviposition day (*X*^2^_1_ = 9248.33, P < 2.2 × 10^−16^), and a significant interaction between both factors (*X*^2^_2_ =30.51, P = 2.4 × 10^−7^). Specifically, the cumulative fecundity in the control lines was lower than the slow-ramping selected lines (Tukey P-adjusted value = 0.02) and the fast-ramping selected (Tukey P-adjusted value = 0.04). Whereas lines from both selection protocols showed similar cumulative fecundity (Tukey P-adjusted value = 0.98). Looking for selection effects on each day (Fig. 3), we observed that females of the selected lines lay more eggs than females of the control lines from day 3 after mating. Additionally, cumulative fecundity was not different between replicate lines (*X*^2^_1_ = 2.03, P = 0.15).

**Figure 3.**
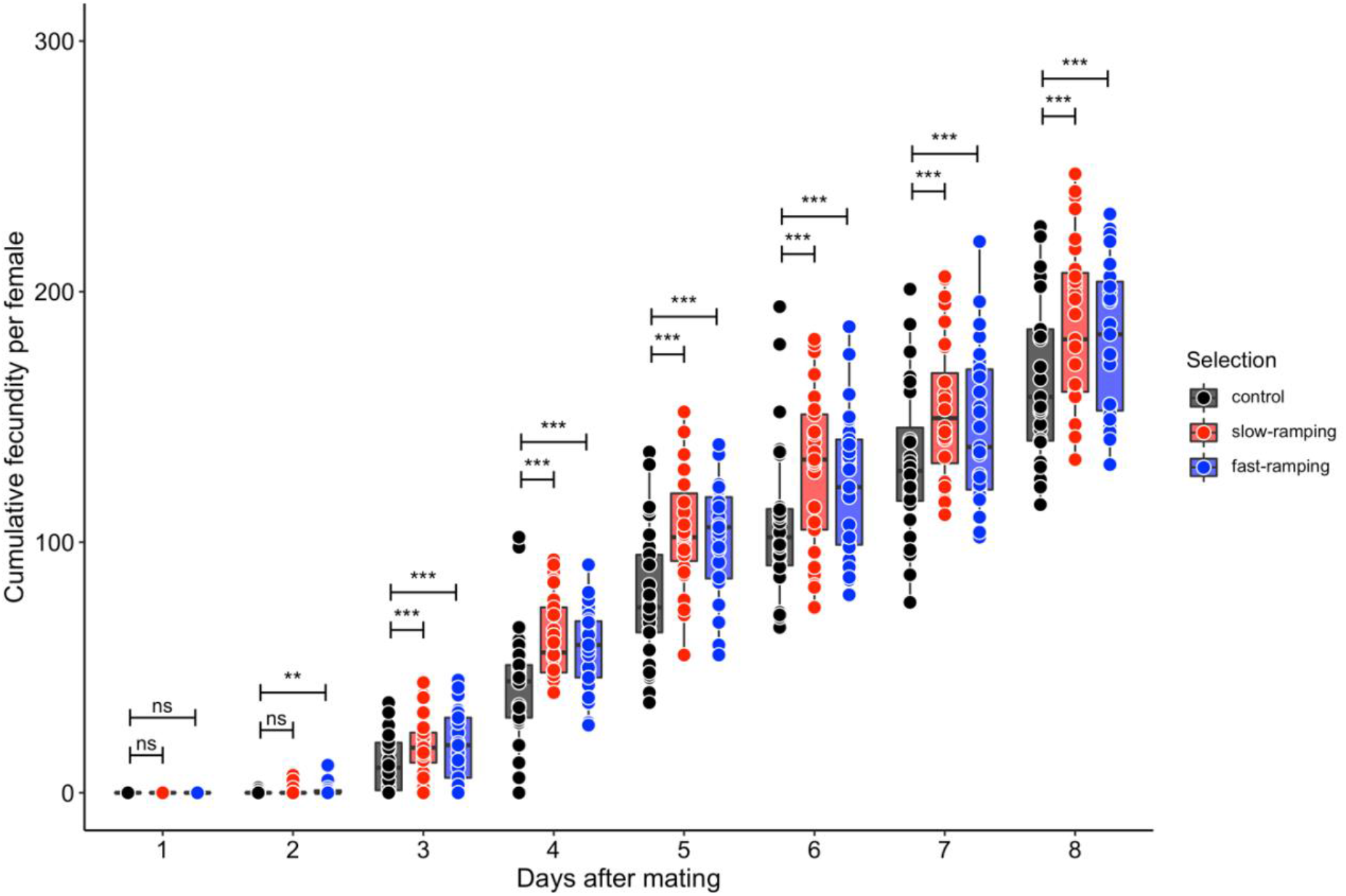
Cumulative fecundity during the first 8 days after mating (early fecundity) for females of control lines (black circles and boxplots), slow-ramping selected lines (red circles and boxplots), and fast-ramping selected lines (blue circles and boxplots) for increasing heat tolerance in *Drosophila subobscura*. Boxplots show the median, the interquartile range (IRQ) and vertical whiskers represent the 1.5*IQR. P-values above horizontal whiskers show the results of a posteriori comparisons between control and selected lines using Tukey tests (ns: P > 0.05, *: P < 0.05, **: P < 0.01, ***: P < 0.001).

For egg-to-adult viability, we found significant effects of thermal selection (*X*^2^_2_ = 16.24, P = 0.0003; Fig. 4). Specifically, slow-ramping selected lines showed a significantly higher egg-to-adult viability than control lines (Tukey P-adjusted value = 0.0002), but not significantly different than fast-ramping selected lines (Bonferroni P-adjusted value = 0.07). On the other hand, fast-ramping selected lines showed similar egg-to-adult viability compared to control lines (Bonferroni P-adjusted value = 0.17). Finally, egg-to-adult viability was significantly different among replicate lines (*X*^2^_4_ = 3.18, P = 1.4 × 10^−6^), but this among-replicates variability did not blur the effects of thermal selection on the egg-to-adult viability.

**Figure 4.**
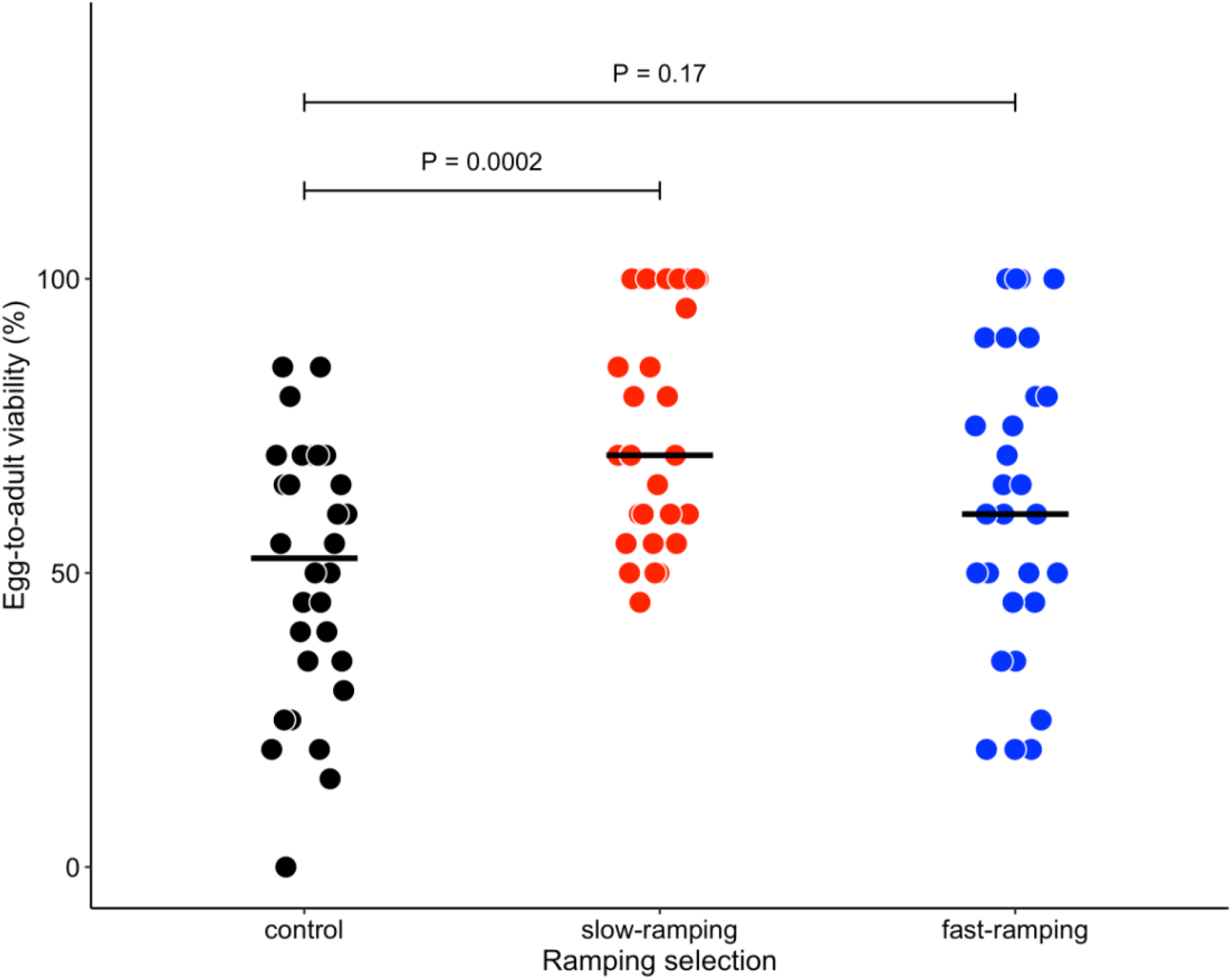
Egg-to-adult viability of flies of control lines (black circles), slow-ramping selected lines (red circles), and fast-ramping selected lines (blue circles) for increasing heat tolerance in *Drosophila subobscura*. Horizontal lines show the mean for each group. P-values above horizontal whiskers show the results of a posteriori comparisons between control and selected lines using Tukey tests.

## Discussion

The mechanisms of tolerance to environmental stress are fundamental for the persistence of natural populations and biological diversity. Temperature is an important abiotic variable that influences the evolution of metabolic and life-history traits in ectotherms (Gilloly et al. 2001; Addo-Bediako et al. 2002; Brown et al. 2004; Padfield et al. 2016, Mallard et al. 2018). In this work, we found that the experimental populations that were selected for an increasing thermal tolerance showed changes in enzyme activities related to energy metabolism, and also evolutionary responses of fitness-related traits in *D. subobscura*. However, the evolution of heat tolerance was not associated with changes in the metabolic rate in selected populations.

Despite the ubiquitous effects of temperature on metabolism, our findings show no evidence of metabolic depression associated with heat tolerance evolution in *D. subobscura*. To explain this finding, we must recognize that there is contrasting evidence about the temperature effects on the evolution of metabolism. For instance, different studies have reported that species living in temperate environments have lower metabolic rates than species inhabiting cold habitats (Addo-Bediako et al. 2002; Sylvestre et al. 2007; Schaefer & Walters, 2010; Sinnatamby et al. 2015), and other studies have found a reduction in metabolic rate in response to warm conditions (Padfield et al. 2016; Mallard et al. 2017; Pilakouta et al. 2020). In line with our findings reported here, there is evidence that does not support the effects of temperature on the evolution of metabolism at intra- (Alton et al. 2017) and interspecific level (Messamah et al. 2017). A plausible explanation is that the evolution of metabolic rate could depend on the interaction between temperature and resource availability, would be difficult to detect metabolic changes under non-stressful maintenance conditions (Alton et al. 2017). A possible explanation could be that, our control and evolved populations were fed ‘ad libitum’ before measures, which could override correlated responses of metabolic rate to heat tolerance evolution. Additionally, metabolic rate was only measured at a single temperature (21°C), which does not allow us to test the effect of heat tolerance evolution on the thermal sensitivity of metabolic rate (e.g., Q_10_). If this is true, it is expected that differences in metabolic rate between selected and control lines would have been larger at higher temperatures (Colinet et al. 2015).

Another explanation for our findings is that metabolic rate is a complex trait that represents the total flux of energy at an organismal level and it does not necessarily account for changes in enzymes related to energy metabolism (O’Brien & Suarez, 2001). For branching pathways as the G6P-branchpoint, there is evidence that selection can acts on enzymes capable of controlling the flux allocation in *Drosophila* species (Flowers et al., 2007). Here, we found evidence that artificial selection for increasing heat tolerance changes the activity of the HEX and G6PD enzymes: HEX evolved to lower enzyme activity in the slow-ramping selected lines, but this effect was only detected when flies were exposed to non-stressful thermal conditions (21°C); whereas the G6PD activity showed significant differences between thermal selection regimens, but the a posteriori analysis did not show differences between evolved and control lines (likely because of the P-value was close to the significance threshold). These changes in the activity of the G6P branchpoint enzymes is concordant with a study that found *D. simulans* populations evolving in thermally fluctuating environments, with a reduction in the gene expression of enzymes involved in energy processing (Mallard et al. 2018). Particularly, we think that the lower enzyme activity could be an evolutionary consequence to heat tolerance evolution in slow-ramping selected lines because flies with high heat tolerance should have had a low metabolism to survive the heat stress for a longer time (Santos et al. 2012).

Our results for the fitness-related traits also provide indirect evidence for the adaptive value of the energy-saving strategies (e.g., metabolic depression). Several studies have found associations between metabolic enzymes and reproductive traits in insects (Kageyama & Ohnishi, 1971; Clark & Fucito, 1998; Harshman et al. 1999; Mallard et al. 2018). We propose that this association can be explained by the pace-to-life syndrome, which proposes that individuals with fast metabolism should grow faster than their counterparts with slow metabolism at the cost to have a shorter lifespan and lower fecundity (Stearns 1989; Polverino et al. 2018; Tüzün et al. 2022). However, our findings only match with the pace-to-life syndrome because a reduced metabolic activity was found only for the slow-ramping selected lines. Whereas an explanation for the higher fecundity exhibited by the fast-ramping selected lines is unclear, despite observing that fast-ramping selected lines exhibited a lower activity of G6PD than control lines (but this difference was not statistically significant). Regarding egg-to-adult viability, slow-ramping selected lines exhibited viability 1.5 times higher than control lines with several vials reaching up to 100% viability, whereas fast-ramping selected lines did not differ from control lines. Previous results have found that low activities of HEX and G6PD were associated with short development times in *D. melanogaster* and *D. subobcura* (Marinkovié et al. 1986), whereas *D. melanogaster* flies that exhibited low activity of HEX (as well as other metabolic enzymes) showed longer lifespan (Talbert et al., 2015).

In conclusion, heat tolerance evolution had positive consequences on fitness-related traits, including increased fecundity and preadult survival. Despite some evidence showing that upper thermal limit (CT_max_) had limited evolutionary potential (Kellermann et al. 2012; Kelly et al. 2012), our research group has found that CT_max_ of *D. subobscura* populations have enough genetic variation (Castañeda et al. 2019), to adapt to local conditions (Castañeda et al. 2015) and respond to artificial selection (Mesas et al. 2021; but see Santos et al. 2022 for no evolutionary response to warm temperatures). Thus, heat tolerance can evolve under different thermal scenarios but with different outcomes on associated traits depending on the intensity of thermal stress. Therefore, spatial and temporal variability of thermal stress intensity should be taken into account in future studies (see Buckley et al., 2013; Rezende et al. 2020) if we want to understand and predict the adaptive response to ongoing and future climatic conditions.

## Acknowledgements

We thank Inês Fragata, Pedro Simões, Marija Savić Veselinović, Ramakrishnan Vasudeva, and one anonymous reviewer for their comments and suggestions during the review process. We also thank Angélica Jaramillo for her valuable help in the fly maintenance and experiments, Roberto Nespolo for facilitating the respirometry equipment, and Kristi Montooth for providing us with the protocols to measure enzyme activities. A.M thanks a CONICYT fellowship no. 21140595 (Chile).

Preprint version 4 of this article has been peer-reviewed and recommended by Peer Community In Evolutionary Biology (https://doi.org/10.24072/pci.evolbiol.100155)

## Data, scripts, code, and supplementary information availability

Data and scripts are available online: https://doi.org/10.6084/m9.figshare.20373180.v1 Supplementary information (Figure S1) is available at the end of the present article.

## Author contributions

A.M. designed experiments, conducted experiments and statistical analyses and wrote the first draft of the manuscript. L.E.C. conceived the original idea, designed experiments, conducted statistical analyses, provided funds for all experiments, and edited and wrote the manuscript.

## Supplementary Material

**Figure S1.**
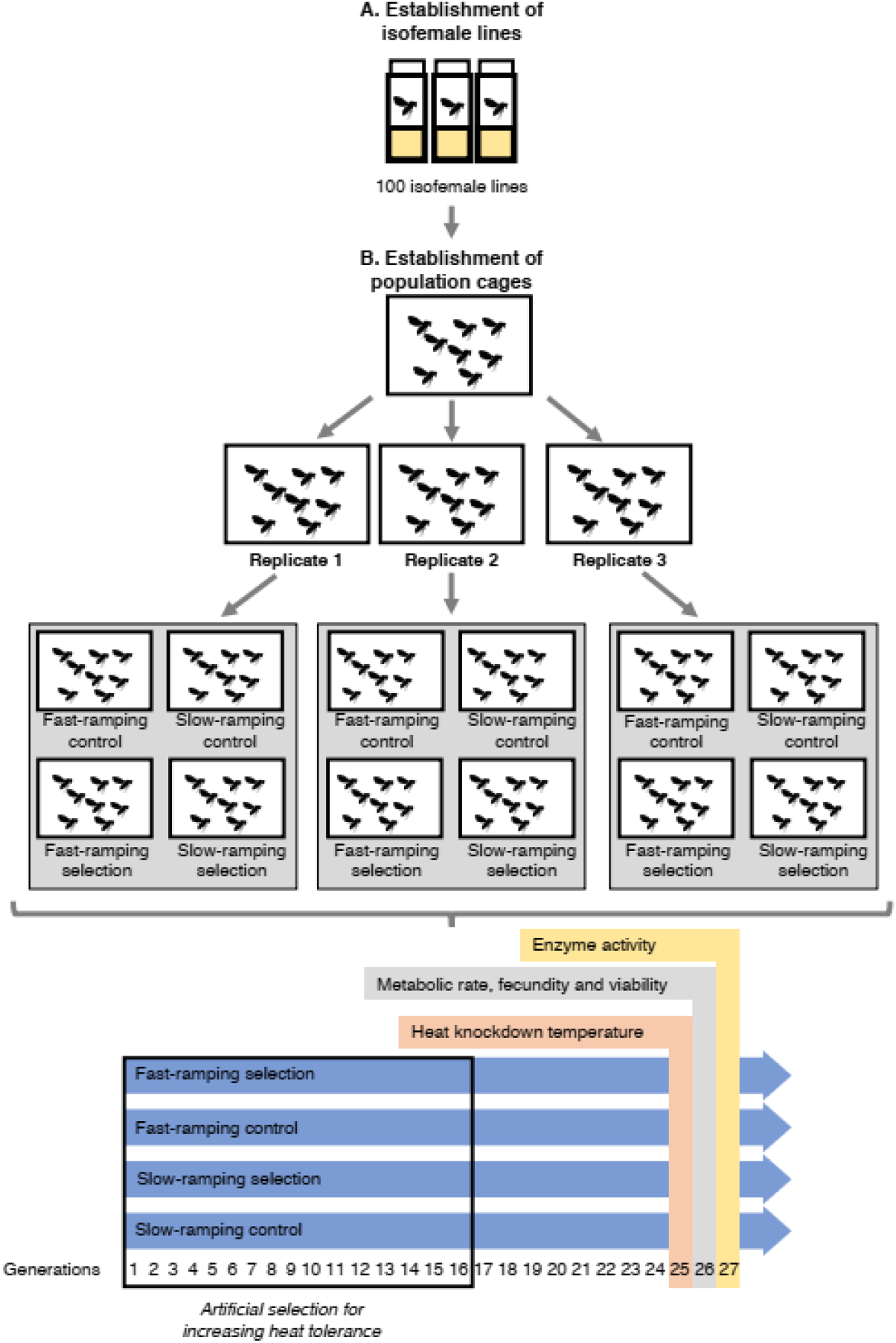
Experimental design: 100 isofemale lines from *Drosophila subobscura* were used to establish an outbred population. The F1 of these isofemale lines were transferred to a population cage and the F2 flies were divided into three replicates: R1, R2, and R3. After 3 generations, each population cage was divided into four population cages, which were assigned to four different artificial selection protocols in triplicate: fast-ramping selection, fast-ramping control, slow-ramping selection, and slow-ramping control lines. During 16 generations, heat tolerance was selected for 33% highest values of knockdown temperature using two different selection protocols: slow ramping rate (0.08 °C/min) and fast ramping rate (0.4 °C/min). Traits were evaluated at generation 25, 26 and 27.

