## Supplemental Figure 1 for "Evolutionary responses of energy metabolism, development, and reproduction to artificial selection for increasing heat tolerance in *Drosophila subobscura*"

**Figure S1.** Experimental design: 100 isofemale lines from Drosophila subobscura were used to establish an outbred population. The F1 of these isofemale lines were transferred to a population cage and the F2 flies were divided into three replicates: R1, R2, and R3. After 3 generations, each population cage was divided into four population cages, which were assigned to four different artificial selection protocols in triplicate: fast-ramping selection, fast-ramping control, slow-ramping selection, and slow-ramping control lines. During 16 generations, heat tolerance was selected for 33% highest values of knockdown temperature using two different selection protocols: slow ramping rate (0.08 ºC/min) and fast ramping rate (0.4 ºC/min). Traits were evaluated at generation 25, 26 and 27.

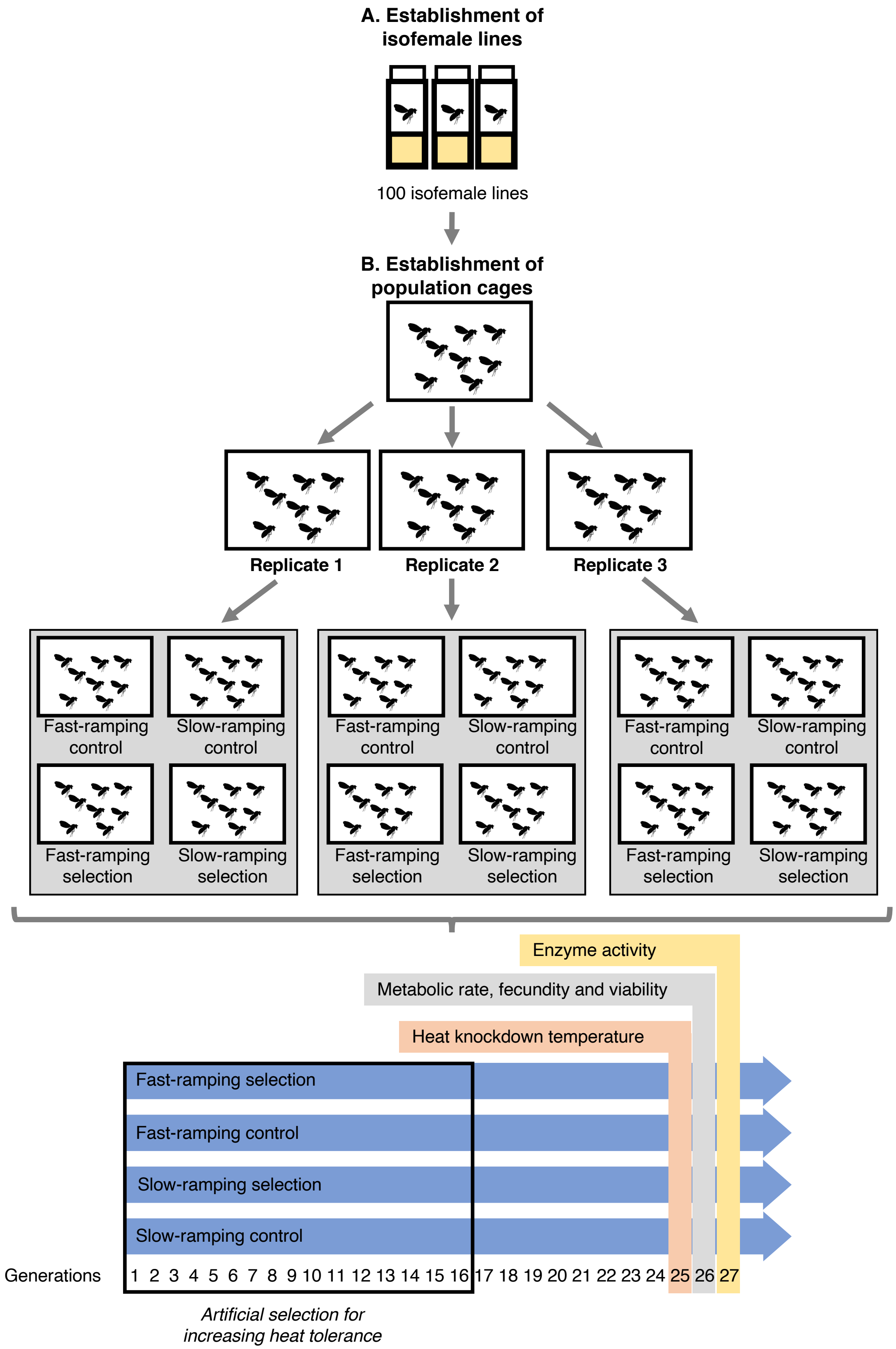
A. Establishment of isofemale lines
100 isofemale lines
B. Establishment of population cages
Replicate 1
Replicate 2
Replicate 3
Fast-ramping control
Slow-ramping control
Fast-ramping selection
Slow-ramping selection
Enzyme activity
Metabolic rate, fecundity and viability
Heat knockdown temperature
Fast-ramping selection
Fast-ramping control
Slow-ramping selection
Slow-ramping control
Generations
1 2 3 4 5 6 7 8 9 10 11 12 13 14 15 16 17 18 19 20 21 22 23 24 25 26 27
Artificial selection for increasing heat tolerance
